# Distributed affective space represents multiple emotion categories across the brain

**DOI:** 10.1101/123521

**Authors:** Heini Saarimäki, Lara Farzaneh Ejtehadian, Enrico Glerean, liro P. Jääskeläinen, Patrik Vuilleumier, Mikko Sams, Lauri Nummenmaa

## Abstract

The functional organization of human emotion systems as well as their neuroanatomical basis and segregation in the brain remains unresolved. Here we used pattern classification and hierarchical clustering to reveal and characterize the organization of discrete emotion categories in the human brain. We induced 14 emotions (6 “basic”, such as fear and anger; and 8 “non-basic”, such as shame and gratitude) and a neutral state in participants using guided mental imagery while their brain activity was measured with functional magnetic resonance imaging (fMRI). Twelve out of 14 emotions could be reliably classified from the fMRI signals. All emotions engaged a multitude of brain areas, primarily in midline cortices including anterior and posterior cingulate and precuneus, in subcortical regions, and in motor regions including cerebellum and premotor cortex. Similarity of subjective emotional experiences was associated with similarity of the corresponding neural activation patterns. We conclude that the emotions included in the study have discrete neural bases characterized by specific, distributed activation patterns in widespread cortical and subcortical circuits, and highlight both overlaps and differences in the locations of these for each emotion. Locally differentiated engagement of these globally shared circuits defines the unique neural fingerprint activity pattern and the corresponding subjective feeling associated with each emotion.

## Introduction

The organization of human emotion systems is currently a topic of lively debate (Hamann, 2012; Lindquist et al., 2012; Kragel and LaBar, 2014; Saarimäki et al., 2016). This discussion revolves around the number of distinct emotion systems and the organization of human emotion circuits in the brain. Most research on discrete emotion categories has focused on ‘primary’ or ‘basic’ emotions (usually anger, fear, disgust, happiness, sadness, and surprise). According to the basic emotion theories, these have been shaped during the evolution to serve distinct survival functions via discrete neural circuits and physiological systems (Ekman, 1992; Ekman et al., 1999; Panksepp, 1982; Damasio 1999). Several human neuroimaging studies support the view that the basic emotions have discrete neural bases, as they are associated with discernible neural activity patterns as measured by BOLD-fMRI (e.g. Kragel and LaBar 2014; Saarimäki et al., 2016; for a review, see Kragel and LaBar 2016).

Yet, a wide array of other emotions, including ‘secondary’ or ‘social’ emotions^1^ (see reviews and proposed taxonomies in Damasio 1999; Adolphs 2002a), also serve adaptive survival functions and are characterized by distinctive facial expressions (Baron-Cohen et al., 2001; Shaw et al., 2005), bodily sensations (Nummenmaa et al., 2014a), and neural activity patterns (Kassam et al. 2013; Kragel and LaBar 2015). Nevertheless, the psychological and neural mechanisms of these non-basic emotions, as well as their commonalities or differences relative to basic emotions, remain largely unresolved (Ekman, 1999; Ekman and Cordaro, 2011; Adolphs, 2002b). In particular, these emotions may involve more elaborate cognitive representations acquired through experience, education, and social norms (Panksepp and Watt, 2011), and hence, recruit brain systems partly distinct from those implicated in more “primitive” (and possibly partly innate) basic emotions. It is thus possible that also non-basic emotions may have discrete neural bases which would, however, be discernible from that of basic emotions.

A set of core emotion processing regions is consistently engaged during multiple emotions. These include cortical midline regions (Peelen et al., 2010; Chikazoe et al., 2014; Trost et al., 2012), somatomotor regions (Adolphs et al., 2000; de Gelder et al., 2004; Pichon et al., 2008; Nummenmaa et al., 2008, 2012), as well as amygdala, brainstem, and thalamus (Adolphs, 2010; Damasio and Carvalho, 2013; Kragel and LaBar, 2014). These regions serve as candidate areas containing discrete neural signatures for different basic emotions (Peelen et al., 2010; Saarimäki et al., 2016). Yet, it is currently unclear whether these regions also code for other emotions. Prior studies using univariate analyses have quantified neural responses to emotions such as regret (Eryilmaz et al., 2011; Coricelli et al., 2005), guilt (Wagner et al., 2011), pride (Takahashi et al., 2008; Simon-Thomas et al., 2012; Zahn et al., 2009), rejoice (Chandrasekhar et al., 2008), and maternal love (Bartels and Zeki, 2004), as well as aesthetic feelings such as wonder or nostalgia (Vuilleumier and Trost, 2015). Yet, studies (except Kassam et al., 2013; Kragel and LaBar 2014) have usually compared brain activation differences between two emotions in a univariate fashion. Thus, it remains unclear whether similar, distinct circuits as previously observed for basic emotions would also support these types of emotions.

Here, we investigated the neural underpinnings of multiple basic and non-basic emotion categories using multivariate pattern recognition (MVPA) and multidimensional scaling. We induced fourteen emotions in participants by guided mental imagery while their brain activity was measured with fMRI. First, to examine whether different emotions have discrete brain bases, we employed pattern classification. Successful pattern classification would thus reveal that specific neural signatures consistently characterize different emotions. Second, using hierarchical clustering, we examined the similarity of neural substrates of different emotions and tested how this similarity was related to how similarly these emotions are experienced. Third, we mapped representation of different emotions in the core emotion processing regions of the brain, by characterizing the emotion-dependent neural response profiles using region-of-interest pattern classification, univariate analyses, and cumulative activation mapping.

## Materials and Methods

### Participants

Twenty-five female volunteers (ages 19–38, mean age 23.6 years) participated in the experiment. All were right-handed, neurologically healthy and with normal or corrected-to-normal vision, and gave written informed consent according to the Declaration of Helsinki. The Institutional Review Board of Aalto University approved the experimental protocol. Female participants were chosen to maximize the power of the experiment, as when compared to males, females typically experience and portray more intensive emotional reactions, and show greater facial mimicry as indexed by EMG (Grossman and Wood, 1993).

### Stimuli

The stimuli for the guided affective imagery were sixty 5–20 second long narratives describing an antecedent event triggering prominently one emotional state. Narrative-based guided imagery is known to be an efficient emotion induction technique that engages widespread brain emotion circuits (Costa et al., 2010; Nummenmaa et al., 2014b), consistent with a reliable impact on affective state. Each narrative elicited primarily one out of possible 14 emotions, or a neutral emotional state. Targeted emotions included six basic or primary emotions (anger, fear, disgust, happiness, sadness, and surprise) and eight social or secondary emotions (shame, pride, longing, guilt, love, contempt, gratitude, and despair). We initially created ten narratives for each category (total of 150 narratives). The narratives included a short description of a situation that participants were instructed to imagine would happen to them, for instance, *“It is a late night on a dimly-lit parking lot. Your car stands alone in a dark corner. Suddenly you hear a gun shot behind you*.” (fear), or *“Your lover is lying next to you on a bed. You look into his eyes when he gently touches your hair and bends to kiss your lips.”* (love). In an online pilot study, 50 participants rated the discrete emotion best corresponding to each narrative (for details, see Supplementary Material). Subsequently, we selected four narratives per category (total of 60 narratives; Supplementary Table S1) that were most consistently labeled with the specific target category to be used in the main experiment.

The selected narratives were spoken by a female speaker using neutral prosody without cues for the affective content of the narrative. The background noise in the recording room was recorded and equalized (bandpass filter 50–10000 Hz) with Apple Logic Pro 9 (Apple Inc.), and gate and compressor were used to attenuate the background noise during moments of silence and slightly compress the voice dynamic to limit the variance of the sound power. The loudness of each narrative was normalized according to the peak value.

The recorded narratives were randomly divided into four runs of fifteen narratives, with one narrative per category in each run. The runs contained the same narratives for all participants but the presentation order was randomized within each run and for each participant. During fMRI, the four sets of recorded narratives were all presented twice in a random order thus resulting in altogether eight runs. Each run lasted 7–8 minutes and consisted of 15 trials. A trial started with a fixation cross shown for 0.5 seconds, followed by a 2-s presentation of a word describing the target emotion (anger, fear, disgust and so forth) to prepare participants for the forthcoming emotional imagery task and thus to make the induction more powerful. Next, the narrative was spoken out, followed by a 10-s imagery period. The trial ended with a 10-s wash-out period to counter for possible carryover effects. Subjects were instructed to listen to the narratives similarly as if they would listen to a radio or podcast, and to try to get involved in the narratives by imagining the described events as happening to themselves and to experience the corresponding feeling as vividly as possible.

Auditory stimuli were delivered through Sensimetrics S14 insert earphones (Sensimetrics Corporation, Malden, Massachusetts, USA). Sound was adjusted for each subject to be loud enough to be heard over the scanner noise. The visual stimuli were delivered using Presentation software (Neurobehavioral Systems Inc., Albany, CA, USA), and they were back-projected on a semitransparent screen using a 3-micromirror data projector (Christie X3, Christie Digital Systems Ltd., Mönchengladbach, Germany), and from there via a mirror to the participant.

After the scanning, participants were presented with the narratives again, and, using an on-line rating tool, rated how strongly they felt each of the possible 14 (and neutral state) emotions using a scale ranging from 0 (not at all) to 9 (very much), which was for further analyses scaled to range from 0 to 1 (for details, see Supplementary Material). The resulting intensity profiles (i.e., intensity ratings per category for each narrative) are shown in Figure 1a. We also ran k-means clustering for these intensity profiles to test whether the 15 target categories could be identified from the emotion-wise intensity ratings, thus revealing whether or not the narratives elicited discrete categorical emotions. Also, to test the similarity of narratives belonging to the same category, we calculated the Euclidean distance between the intensity profiles of each pair of narratives (Figure 1b).

**Figure 1.**
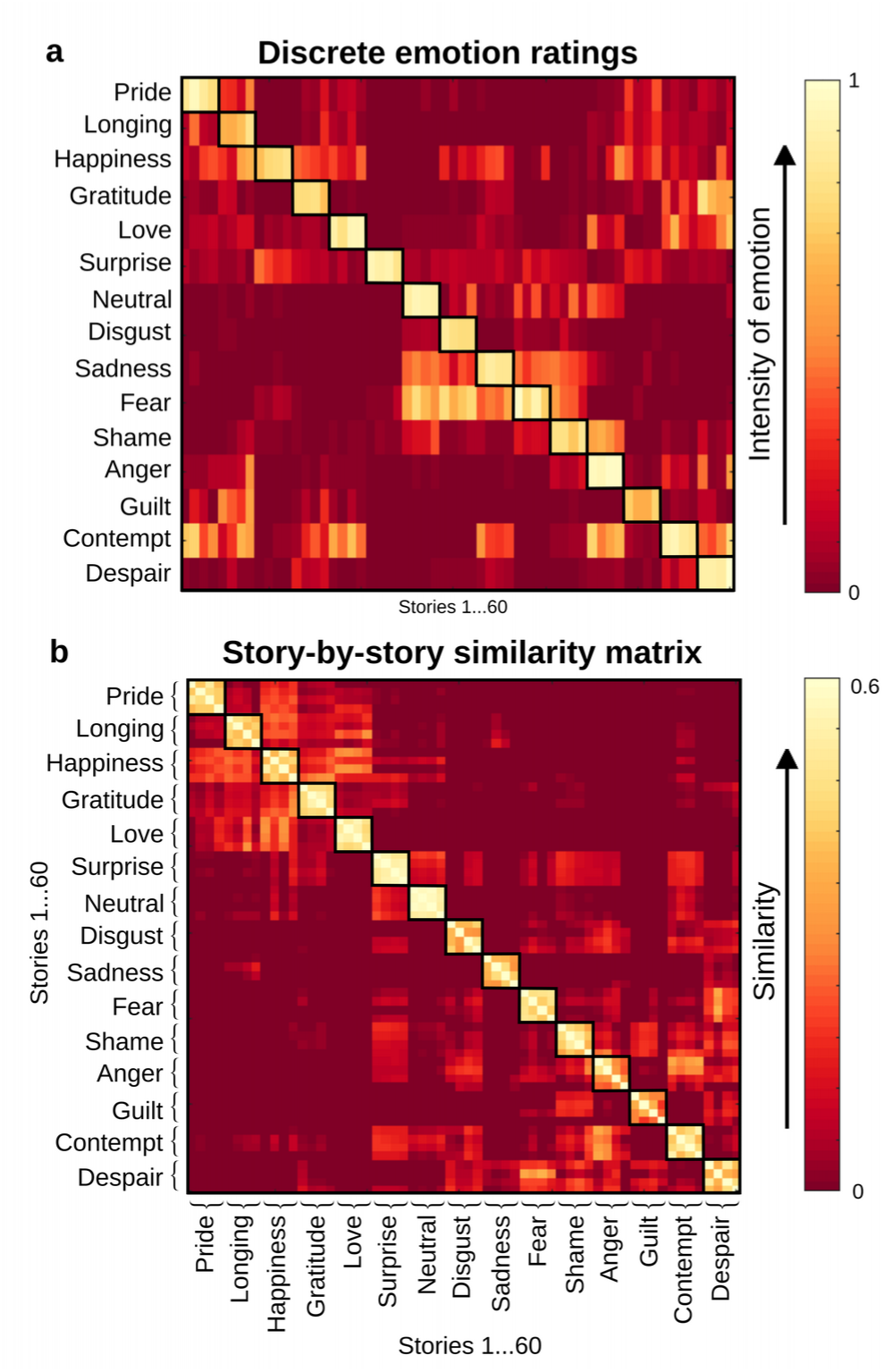
The stimuli consisted of 60 brief (5–20 s) narratives that induced 14 emotional states and a neutral state. a) Participants rated on a scale from 0 to 1 how much of each emotion was elicited by the narrative. The coloring indexes the mean intensity for experiencing each emotion for each narrative. b) Based on the ratings, we calculated the similarity of emotion content between narratives by using Euclidean distances.

Finally, participants rated the similarity of all possible pairs of the emotions induced during the experiment (a total of 105 pairs) using a direct comparison method for the emotion words corresponding to the target emotions. The participants were shown one pair of emotion words at the time and asked to rate the similarity between the emotion pair (ranging from no similarity [0] to full similarity [5]). The ratings were then scaled to range between 0 and 1 and used to extract the experiential similarity matrix first for each participant individually and these were then averaged over all participants.

### MRI data acquisition and preprocessing

MRI data were collected on a 3T Siemens Magnetom Skyra scanner at the Advanced Magnetic Imaging center, Aalto University, using a 20-channel Siemens volume coil. Whole-brain functional scans were collected using a whole brain T2*-weighted EPI sequence with the following parameters: 33 axial slices, TR = 1.7 s, TE = 24 ms, flip angle = 70°, voxel size = 3.1 × 3.1 × 4.0 mm^3^, matrix size = 64 × 64 × 33, FOV 256 × 256 mm. A custom-modified bipolar water excitation radio frequency (RF) pulse was used to avoid signal from fat. High-resolution anatomical images with isotropic 1 × 1 × 1 mm^3^ voxel size were collected using a T1-weighted MP-RAGE sequence.

Data were preprocessed using FSL 5.0 (Jenkinson et al., 2012). Motion correction was performed using MCFLIRT (Jenkinson et al., 2002) and non-brain matter was removed using BET (Smith, 2002). High-pass temporal filtering was applied using Gaussian-weighted least-squares straight line fitting with sigma of 100s. Participant-wise gray matter masks were generated by segmenting the T1-weighted images into gray and white matter and cerebrospinal fluid using the FAST segmentation tool (Zhang et al., 2001) and transforming the masks to the native space using FLIRT (Jenkinson et al., 2002) with 9 degrees of freedom. The gray matter maps were subsequently thresholded using intensity threshold > 0.5 to create participant-specific masks. This threshold was chosen to include those voxels with a higher probability of belonging to the gray matter and, subsequently, the masks were visually inspected to make sure most gray matter was included. On average, the gray matter mask included 16000 voxels.

For univariate GLM analysis, the preprocessed functional data were registered to 2-mm Montreal Neurological Institute (MNI) 152 standard space template using FLIRT (Jenkinson et al. 2002). The brain-extracted T1-weighted images were first normalized to the MNI space and the normalization parameters were subsequently applied to the EPI images. All registrations were performed using 9 degrees of freedom. Finally, the functional data were smoothed by using a Gaussian kernel with FMWH 8.

### Multivariate pattern classification within participants

The classification of emotion categories was performed using the Princeton multi-voxel pattern analysis (MVPA) toolbox (https://pni.princeton.edu/pni-software-tools/mvpa-toolbox) in Matlab 2012b using each participant’s data in native space. A separate classifier with the following pipeline was trained for each participant and, after all steps, the classification results were averaged across the participants. After the preprocessing steps described in the previous section, for each participant’s fMRI data, voxels outside gray matter were masked out using the participant-specific gray matter masks and the functional data from each run were standardized to have a mean of zero and variance of one. Next, each participant’s data were divided into training (N-1 runs) and testing sets (the remaining run). Feature selection was performed using one-way ANOVA (testing for the main effect of emotion category) for the training set to select the voxels with a significant (p<0.05) main effect for emotion, i.e., to select the voxels whose mean activation differed between at least some of the 15 possible emotion conditions. The feature selection preserved on average (across cross-validation folds and participants) 41% of the voxels. Haemodynamic lag was corrected for by convolving the boxcar category regressors with the canonical double gamma HRF function and thresholding the convolved regressors using a sigmoid function to return the regressors to the binary form. The classification was performed on all the standardized, HRF-convolved fMRI volumes from the 10s imagery period following the narrative (treating all single time points per category as samples for that category; median 6 volumes per one emotion category in one run) to extract only emotional effects, and to minimize activity related to the acoustic and semantic features of the stimuli. Thus, each of the eight runs included on average 90 TRs of stimulation (5-6 TRs per category) that were used in the classification. A linear neural network classifier without hidden layers was trained to recognize the correct emotion category out of 15 possible ones (multiclass classification, see Polyn et al. 2005 for details). Naïve chance level, derived as a ratio of 1 over the number of categories, was 6.7%. A leave-one-run-out cross-validation was performed, thus, dividing the data into all possible N-1 run combinations and repeating the classification pipeline for each such cross-validation fold, and the participant-wise classification accuracy was calculated as an average percentage of correct guesses across all the cross-validation folds. To test whether classification accuracy exceeded chance level, we used permutation tests to simulate the probability distribution of the classification. Each permutation step included shuffling of category labels of the training set (across training set runs) and re-running the whole classification pipeline, repeated 1,000 times for each subject. FDR correction at p<.05 was used for multiple comparisons.

### Region-of-interest analysis

We also applied a region-of-interest (ROI) analysis to test whether the classification accuracies of emotions differ in any of our *a priori* defined regions of interest. Cortical regions showing consistent emotion-related activation in the literature were selected as candidate ROIs for coding discrete emotional content (Kober et al., 2008; Vytal and Hamann, 2010): orbitofrontal cortex (OFC), frontal pole, inferior frontal gyrus (IFG), insula, anterior cingulate cortex (ACC), posterior cingulate cortex (PCC), medial frontal cortex (MFC), precuneus, paracingulate gyrus, precentral gyrus, supplementary motor area, and postcentral gyrus. The subcortical regions were amygdala, nucleus accumbens, hippocampus, and thalamus. Bilateral masks for these ROIs were first defined in MNI standard space using the Harvard-Oxford cortical and subcortical atlases (Desikan et al., 2006) and then transformed into native space using FLIRT in FSL. Feature selection and classifier training was then performed for each ROI separately similarly to the whole brain analyses.

### Hierarchical clustering

We next investigated the similarities between emotions using the category confusions from the whole-brain classification. From the group-averaged confusion matrix, we calculated a distance matrix by taking the category confusion vectors for each pair of emotions and by calculating the Euclidean distance between these vectors (see Reyes-Vargas et al., 2013). We then employed hierarchical cluster analysis in Matlab to investigate how different emotions cluster together based on their neural similarities.

The agglomerative hierarchical cluster tree was calculated on the distance matrix using *linkage* function with *‘complete’* option (i.e., the furthest distance method). Finally, we constructed the clusters from the cluster tree (*cluster* function) and chose the clustering that minimized the variance of the number of elements in the clusters. To visualize the similarities in subjective and neural organization of emotions, we extracted the clusters in both neural and behavioral data, and subsequently plotted the cluster solutions using alluvial diagrams (Rosvall and Bergstrom, 2010; www.mapequation.org).

Finally, we investigated to which extent the neural similarities between different emotional states correspond to their experiential (subjectively felt) differences. Experiential similarity matrices were calculated based on pairwise similarity ratings of emotions and averaged over the participants. Subsequently, the mean neural and subjective similarity matrices were correlated using Spearman’s rank correlation coefficient. The *p* level for the Spearman test was obtained with a permutation test by shuffling the neural matrix and re-calculating the correlation for 10^^^6 times.

### Regional effects in GLM

To investigate the overall effect of any emotion on brain activity, we first ran general linear model (GLM) to compare all emotions together against the neutral baseline. First level analysis was performed in SPM 12 (wwwl.fil.ion.ucl.ac.uk/spm/) to obtain individual contrast maps and second level analysis was then performed with FSL randomise with the threshold free cluster enhancement (TFCE) option as it implements the least biased way to control for multiple comparisons (Eklund et al. 2016; N=5000 permutations).

Next, we ran separate GLMs to compare each of the 14 emotions versus the neutral baseline. For each emotion we computed first level analysis and obtained individual contrast maps. Second level analysis was then performed with FSL *randomise* with TFCE (N=5000 permutations). Results were then qualitatively summarized across emotions by visualizing a cumulative map where each voxel shows the number of statistically significant emotions, at the cluster corrected level of p < 0.05.

To quantify the effect of the findings, we also computed average effect sizes across emotions by first converting unthresholded *t* values to R values for each emotion and averaging them with the Fisher’s formula. Confidence intervals for R values were obtained with bootstrapping. Average effect size (see Supplementary Figure S4) and cumulative maps resulted to be very similar, with a spatial overlap of r=0.7425 (overlap computed with Pearson’s correlation, original *t* maps are available in http://neurovault.org/collections/TWZSVODU).

### Visualization of emotion clusters in the brain

Finally, to summarize and visualize where emotion-wise activation patterns were located, we mapped the three principal clusters obtained with hierarchical clustering on the cortical and subcortical maps using R, G, B color space. The last cluster containing surprise and neutral was plotted separately (see Supplementary Figure S6) given it contained neutral-like states only. For each emotion, we took the unthresholded second level *t* maps obtained from the GLM analysis, summed them for emotions belonging to the same cluster, and assigned the summed values to the corresponding R, G, B channels. The color channels were subsequently visualized in MNI space. Consequently, the RGB color at each voxel reflects the cluster distribution of that voxel, and can be used for localizing brain regions contributing to different emotions.

## Results

### Behavioral results

Behavioral ratings showed that the narratives successfully elicited reliable and strong target emotions (intensity profiles in Figure 1a; mean intensities per target category: pride 0.80, longing for 0.77, happiness 0.84, gratitude 0.84, love 0.90, surprise 0.76, neutral 0.88, disgust 0.85, sadness 0.96, fear 0.89, shame 0.75, anger 0.68, guilt 0.91, contempt 0.66, despair 0.87). In k-means clustering, the accuracy to assign a narrative to the correct target category based on its intensity profile was 97% (against the chance level 6.7%). Also, the narratives within each category had highly similar intensity profiles (Figure 1b); i.e., narratives belonging to the same category elicited similar emotions.

### Classification of basic and non-basic emotions

To test whether different emotions have discernible neural signatures, we trained a linear neural network classifier without hidden layers to recognize the corresponding emotion category from whole-brain fMRI data. Mean classification accuracy across the 14 emotions and the neutral state was 17% (against naïve chance level of 6.7%; 95^th^ percentile of the permuted classification accuracy distribution was 8.4%). After correcting for multiple comparisons, the classification performance was above a permutation-based significance level for all emotions except shame and longing for (*p* < 0.05, Figure 2; see Supplementary Table S2 for effect sizes and Supplementary Figure S1 for a confusion matrix). On average, basic emotions (anger, disgust, fear, happiness, sadness, surprise) could be classified more accurately than the social emotions (26% vs. 15%, respectively, t(24)=7.39, p <0.0001, Cohen’s h=0.16). To examine how emotions could be classified in different single brain regions, we trained a classifier for each of our *a priori* defined regions-of-interest in the emotion circuit separately (see Methods). Classification accuracies for each emotion, organized based on the clustering of whole-brain classifier confusions (see Figure 4), in each region of interest are shown in Figure 3 (see also Supplementary Table S3). In general, classification accuracies were above chance level in frontal ROIs, especially in frontal pole, and in somatomotor ROIs, especially in pre- and postcentral gyri. We also compared the mean classification accuracies of basic and non-basic emotions (Figure S2). Mean classification accuracy exceeded chance level (p<0.05) for both basic and non-basic emotions in frontal pole, precentral gyrus, paracingulate cortex, medial frontal cortex, postcentral gyrus, orbitofrontal cortex, and precuneus, and for basic emotions only in inferior frontal gyrus, posterior and anterior cingulate, insula and supplementary motor area. No ROI showed successful classification for non-basic emotions only. In general, classification accuracies were slightly higher for basic emotions; however, a statistically significant difference between classification accuracy for basic vs. non-basic emotions was found only in frontal pole (t(24)=3.4, p=0.04).

**Figure 2.**
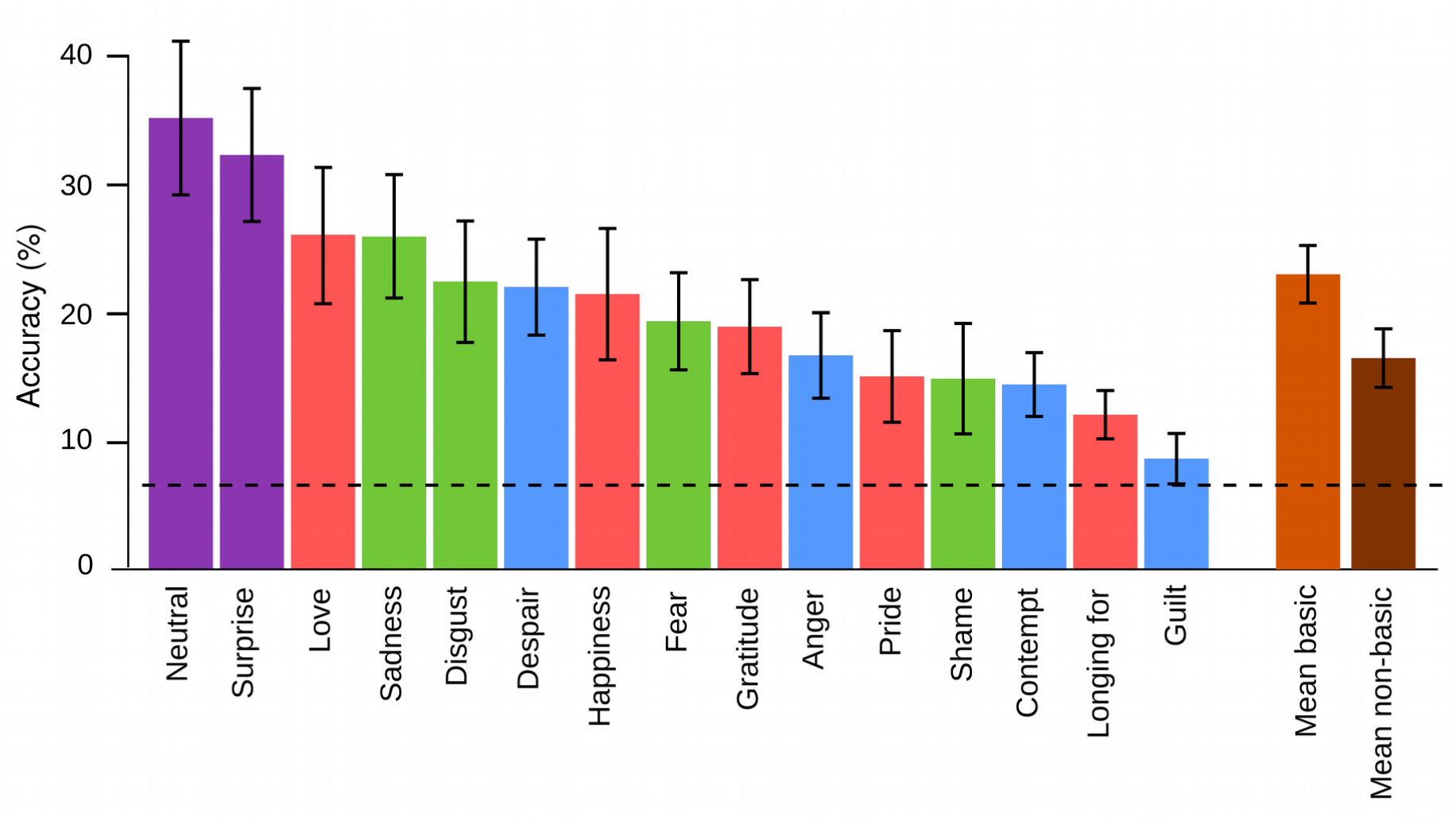
Means and standard errors for emotion-wise whole-brain classification accuracy. Dashed line represents chance level (6.7%). Colors reflect the clusters formed on the basis of experienced similarity of emotions (see Figure 4).

**Figure 3.**
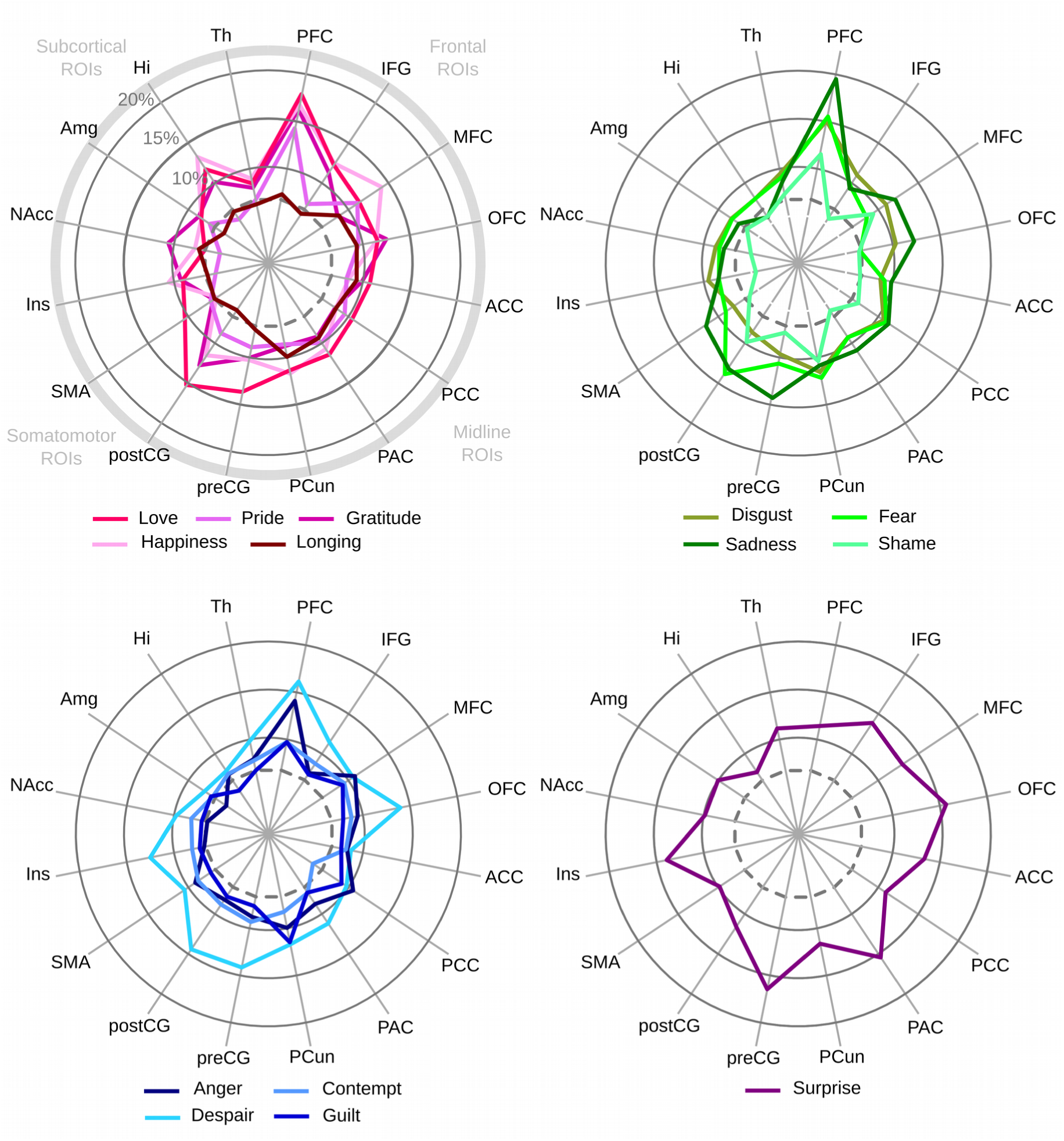
Region-wise classification accuracies for each emotion. Dashed line represents the chance level (6.7%). The colors and clustering is based on the cluster solution shown in Figure 4. PFC = frontal lobe, IFG = inferior frontal gyrus, MFC = medial frontal cortex, OFC = orbitofrontal cortex, ACC = anterior cingulate cortex, PCC = posterior cingulate cortex, PAC = paracingulate cortex, PCun = precuneus, preCG = precentral gyrus, postCG = postcentral gyrus, SMA = supplementary motor area, Ins = insula, NAcc = nucleus accumbens, Amg = amygdala, Hi = hippocampus, Th = thalamus.

**Figure 4.**
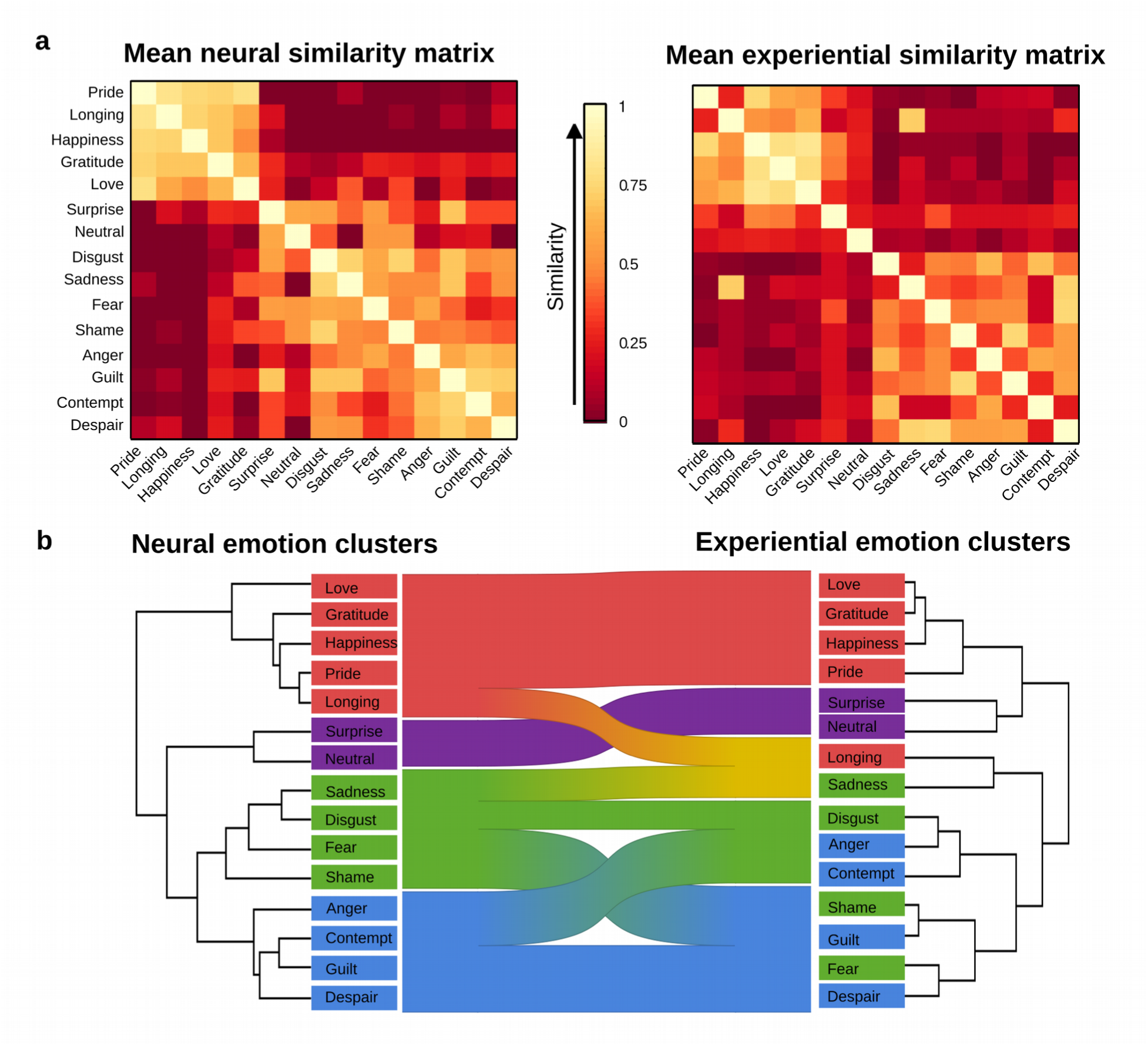
Left: Neural similarity matrix extracted from the classifier confusion matrix. The similarity matrix was created by calculating the Euclidean distance between each pair of emotions based on their category confusion vectors. Right: Experiential similarity matrix based on pairwise similarity ratings for emotions elicited by the narratives. b) Alluvial diagram showing the similarity of hierarchical cluster structure of the experiential and neural similarities. Coloring of the emotion categories is based on the clusters in the neural similarity matrix.

### Similarity of neural basis of different emotions

We then examined the similarity of neural substrates between different emotions using hierarchical clustering on the whole-brain classifier confusions. Clustering of confusion matrices divided the emotions into four clusters (Figure 4a): 1) happiness, pride, gratitude, love, and longing; 2) disgust, sadness, fear, and shame; 3) anger, contempt, guilt, and despair; and 4) surprise and neutral. Next, we investigated the concordance between brain activation patterns and subjective feelings of different emotions with representational similarity analysis. Subjective and neural similarity matrices were averaged across subjects, and compared with Spearman’s rank correlation coefficient. This revealed that organization of subjectively reported similarity of emotions was significantly associated with the similarity of their neural signatures (r=0.68, p<0.0001).

This correspondence was visualized using hierarchical cluster analysis and alluvial diagrams (Figure 4b).

### Affective space in the brain

To investigate the brain regions generally activated and deactivated by our emotion stimuli, we contrasted the brain activity related to all emotions with the neutral condition (Figure 5 outlines; see also Supplementary Figure S3). The areas activated by emotion in general included premotor cortex, thalamus, insula, and putamen. Deactivations were observed in visual and auditory cortices, precuneus, PCC, right anterior PFC, and right lateral parietal areas.

To reveal the brain regions contributing most consistently to different emotions, we next constructed cumulative activation and deactivation maps of emotion-driven haemodynamic responses (Figure 5; the emotion-wise statistical *t* maps are available in http://neurovault.org/collections/TWZSVODU). These maps reveal how many of the 14 possible emotional states activated each voxel. The cumulative activation map (Figure 5a) showed that most emotions involved activation of midline regions including anterior cingulate (ACC) and precuneus, as well as subcortical regions including brain stem and hippocampus, motor areas including cerebellum, and visual cortex. The cumulative deactivation map (Figure 5b) showed that most emotions involved deactivation of auditory cortex, frontal areas including superior frontal gyri, right middle frontal gyrus, and left inferior frontal gyrus, and parietal areas including supramarginal gyrus.

**Figure 5.**
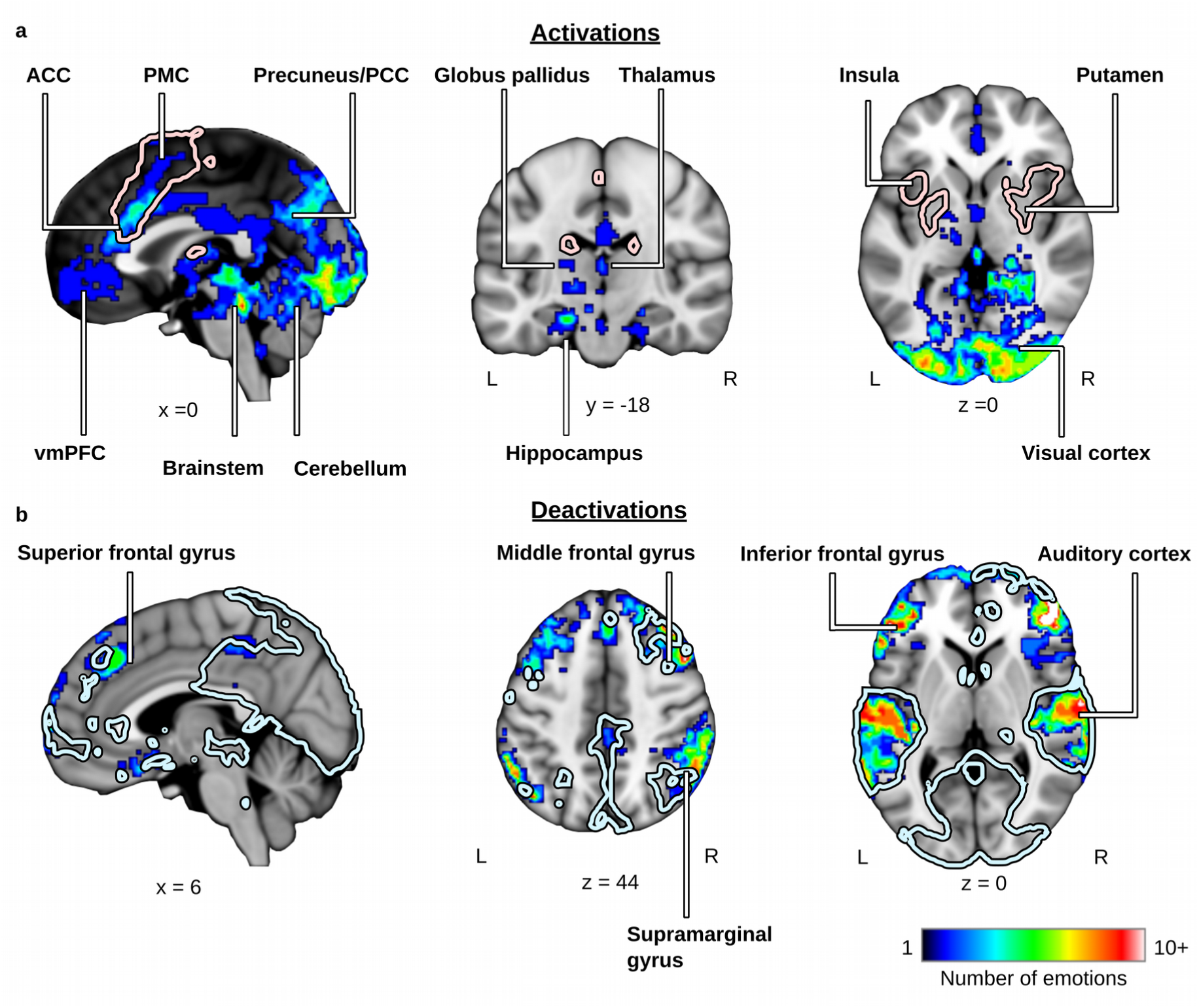
a) Cumulative activation map showing the cumulative sum of binarized *t* maps (p<0.05, cluster-corrected) across each emotion vs. neutral condition. Outline shows the GLM results for all emotions contrasted against the neutral condition p<0.05, cluster-corrected). b) Cumulative deactivation map showing the cumulative sum of binarized *t* maps (p<0.05, cluster-corrected) across neutral vs. each emotion. Outline shows the GLM results for all emotions contrasted against the neutral condition (p<0.05, cluster-corrected).

Finally, to visualize where specific emotions are processed in the brain, we mapped the clusters resulting from the hierarchical clustering on cortical and subcortical surfaces (Figure 6). All emotions activated areas in the visual cortex, ACC, right temporal pole, supplementary motor area, and subcortical regions. In addition, positive emotions belonging to Cluster 1 (happiness, pride, gratitude, love, longing) were more prominent in anterior frontal areas including vmPFC. Negative, self-oriented emotions from Cluster 2 (disgust, sadness, fear, shame) activated especially insula, supplementary motor area, and specific parts of subcortical structures. Negative, other-oriented emotions belonging to Cluster 3 (anger, contempt, guilt, despair; see Supplementary Table S1 for the story stimuli targeting different emotions) were most prominent in left insula and the adjacent frontal areas. Finally, surprise (Cluster 4, Supplementary Figure S6) activated especially parts of auditory cortex, supplementary motor areas, and left insula.

**Figure 6.**
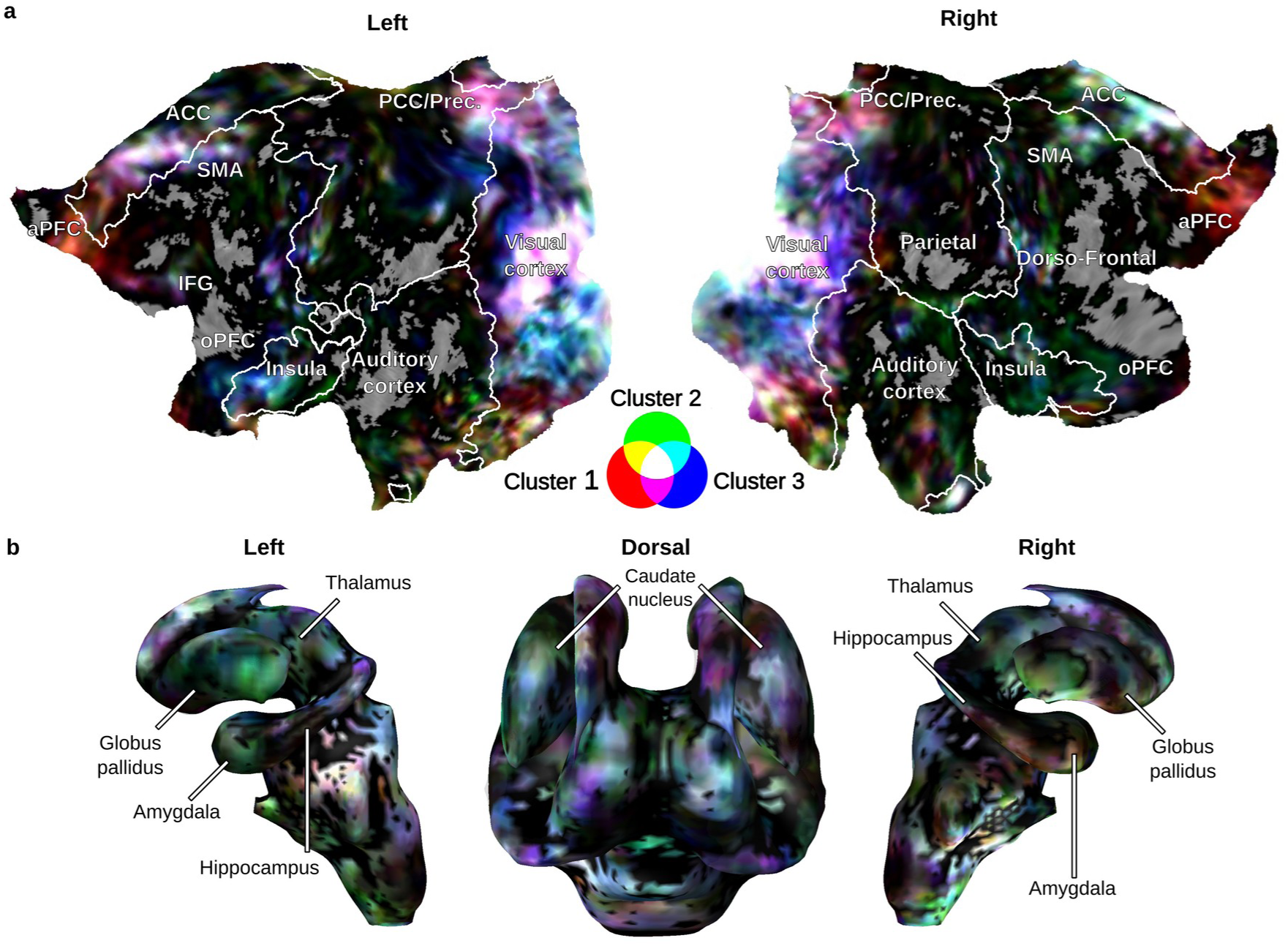
Activation maps showing the summed uncorrected *t* maps for each cluster obtained from the hierarchical clustering analysis in a) cortical regions and b) subcortical regions. Colors represent the three clusters: positive (red), negative self-oriented (green), and negative other-oriented (blue) emotions.

## Discussion

Our results reveal that multiple emotion states have distinct and distributed neural bases, evidenced by the above chance-level classifier performance for all emotions, except longing for and shame. Together with the spatial distribution of emotion-dependent brain activation, this suggests that a multitude of different emotions are represented in the brain in a discrete manner, yet in partly overlapping regions: each emotion state likely modulates the different functional systems of the brain differently, as shown by distinct patterns measured with BOLD-fMRI, and the overall configuration of the regional activation patterns defines the resulting emotion. While a set of ‘core’ emotion processing areas in cortical midline regions, motor areas, sensory areas, and subcortical regions are engaged during practically all emotions, the relative engagement of these areas varies between emotions. This unique neural signature of each emotion might relate to the corresponding subjective feeling, as evidenced by the correspondence between neural and experiential similarity between emotions.

### Different emotions are characterized by discrete neural signatures

Altogether 12 emotions (excluding longing for and shame) out of the 14 included in the study could be reliably classified from the fMRI signals. Our results extend previous studies, which have shown classification of discrete emotional states usually focusing on the basic emotions only or a subset of these (Peelen et al., 2010; Said et al., 2010; Ethofer et al., 2009; Kotz et al., 2012; for a review, see Kragel and LaBar, 2014). While the ‘classic’ basic emotions have attracted most attention in psychological and neurophysiological studies, they constitute only a small portion of the emotions humans universally experience (Edelstein and Shaver, 2007). Furthermore, accumulating behavioral evidence suggests that also other emotions are characterized by distinctive features in facial expressions (Baron-Cohen et al., 2001; Shaw et al., 2005), bodily changes (Nummenmaa et al., 2014a), and physiology (Kreibig, 2010; Kragel and LaBar, 2013). The present data corroborate these findings by showing that also emotions not considered as ‘basic’ may have distinct brain activation patterns. The local brain activity patterns underlying different emotions are most probably to some extend variable across participants and reflect individual responses, as brains are always intrinsically shaped by individual development and experiences.

If we consider discrete emotion systems as wide-spread, discrete neural activation distinct to each emotion state, successful pattern classification of brain states across emotions would provide support for discrete emotion systems. Each emotion likely modulates multiple functional systems of the brain differently, and their spatially distributed configuration might define the specific emotion. For instance, two emotions might share their somatosensory representations, but underlying interoceptive representations could be different. Thus, the general configuration of the nervous system leads to a discrete emotion. However, it must be noted that this type of analysis does not readily reveal the actual neural organization of each emotion system, as the pattern classification only tells us that, on average and at the level measurable with BOLD-fMRI, the pattern underlying each category differ enough to be separated, whereas localizing the actual source of differences is more difficult. For this reason, we have complemented the pattern classification analysis with visualization of different emotion categories using GLM and clustering.

If basic emotions were somehow ‘special’ or different from non-basic emotions at the neural level, we should observe (i) discrete neural activation patterns for basic emotions but not for non-basic emotions, or (ii) different (or perhaps additional) neural systems underlying basic and non-basic emotions. Our classification results and cumulative maps show that both basic and non-basic emotions could be classified accurately to a similar degree, and they elicited activation in largely overlapping brain areas. Further, clustering of category confusions did not provide evidence for a sharp division between basic and non-basic emotions (Figure 4). Nevertheless, classification accuracies within single regions showed that only basic emotions could be distinguished in some areas, including somatomotor regions (insula and supplementary motor area), midline areas (posterior and anterior cingulate), and inferior frontal gyrus, suggesting that on average, these regions might represent basic emotions more accurately than non-basic emotions. Further, the inter-subject classification accuracies (see Supplementary Materials), even though weak for all emotions, were above chance level for basic but not for non-basic emotions, suggesting that neural representations of basic emotions are more shared between individuals than those of non-basic emotions.

### Correspondence between neural and phenomenological organization of emotions

Hierarchical clustering of emotion-specific neural patterns did not follow a clear division between basic and non-basic emotions. The four clusters in the neural data roughly correspond to positive emotions (pride, longing, happiness, gratitude, love), self-oriented negative emotions (disgust, sadness, fear, shame), and other-oriented negative emotions (anger, guilt, contempt, despair), and surprise. Comparison of subjectively experienced similarity of emotions and the similarity of the neural patterns suggested a direct link between the whole-brain neural signatures of emotions and the corresponding subjective feelings: the more similar neural signatures two emotions had, the more similar they were experienced. This accords with prior work suggesting that emotion-specific neural activation patterns might explain why each emotion feels subjectively different (Damasio et al., 2000; Saarimäki et al., 2016). Emotions might constitute discrete activity patterns in regions processing different emotion-related information, such as somatosensory (bodily sensations), motor (actions), as well as brainstem and thalamocortical loops (physiological arousal). Activation from these areas is then integrated in the cortical midline, such integration then giving rise to the interpretation of the subjective feeling (Northoff and Bermpohl, 2004; Northoff et al., 2006). Thus, a subjective feeling of a specific emotion stems from the net activation of different sub-processes, rather than solely on the basis of any single component of emotion.

### Organization of the affective space in the brain

Our hierarchical clustering analysis, cumulative mapping, and region-of-interest classification show in more detail how different patterns of activity may give rise to different emotions. First, midline regions including ACC, PCC, and precuneus were activated during most emotions. Previous research has linked these regions to emotional processing (Lindquist et al., 2012; Phan et al., 2002) and, especially, to the coding of emotional valence (Chikazoe et al., 2014; Colibazzi et al., 2010) but also self-relevance and introspection (Northoff and Bermpohl, 2004; Northoff et al., 2006). These areas are strongly connected to the visceral and subcortical regions (Kober et al., 2008), and are thus thought to integrate information of internal, mental, and bodily states and to hold self-related representations (Buckner and Carroll, 2007; Northoff et al., 2006). Similarly, we found consistent emotion-dependent activity in the brainstem, including periaqueductal grey (PAG), pons, and medulla, for almost all emotions (see Damasio et al., 2000; Damasio, 2010). This activation might reflect the control of autonomic nervous system’s reactions to different emotions (Critchley et al., 2005; Linnman et al., 2012) and/or covert activation of particular motor programs (Blakemore et al., 2016). Also, subcortical regions including amygdala and thalamus showed distinct activity patterns that differed between clusters. Both of these regions are related to salience processing and emotional arousal (Adolphs, 2010; Anders et al., 2004; Damasio and Carvalho, 2013; Kragel and LaBar, 2014) and show specific activation patterns for basic emotions (Wang et al., 2014), findings that we now extent also to nonbasic emotions.

Second, somatomotor areas including premotor cortex, cerebellum (including vermis and the anterior lobe), globus pallidus, caudate nucleus, and posterior insula were activated during most emotions, but according to the cluster visualizations especially during the processing of emotions that have a strong impact on action tendencies and avoidance-oriented behaviors (fear, disgust, sadness, shame, surprise; Frijda et al. 1989). These areas are engaged during emotion perception (Pichon et al., 2008; Nummenmaa et al., 2008, 2012) and emotion regulation (Schutter and van Honk, 2009). They might play a central role in somatomotor processing and action preparation processes related to emotions (Kohler et al., 2002; Wicker et al., 2003; Mazzola et al., 2013).

Third, anterior prefrontal cortex was activated especially during positive emotions (happiness, love, pride, gratitude, longing) according with previous research linking anterior prefrontal cortex with positive affect (Vytal and Hamann, 2010; Bartels and Zeki, 2004; Zahn et al., 2009). Fourth, negative emotions such as guilt, contempt, anger, and despair clustered together, potentially reflecting their social dimension and interpersonal aspects or their self-conscious nature. Especially, left hemisphere activation in orbitofrontal cortex connected to rewards and punishments (Kringelbach and Rolls, 2004), as well as in inferior frontal cortex and dorsolateral prefrontal cortex, which subserve language and executive control (Poldrack et al., 1999; Kane and Engle, 2002), and in anterior insula linked to processing of social emotions (Lamm & Singer, 2010) was activated during these emotions. Fifth, surprise did not resemble any of the other emotions included in this study, but was instead closest to the neutral state. This is in line with previous research showing that surprise tends to separate from other emotions in subjective ratings (Toivonen et al., 2012).

Finally, we also found decreased activation in auditory areas and increased activation in visual areas during the imagery of all emotion categories, likely reflecting the offset of the auditory stimulation followed by mental imagery of the emotion-evoking situation (Ganis et al., 2004) and emotion-related modulation of this activity (Nummenmaa et al., 2012; Holmes and Mathews, 2005; Kassam et al., 2013).

### Limitations

Despite the careful selection of the emotional stimuli and k-means clustering suggesting clear categorical structure in the evoked affect (see Figure 1), it is possible that the narratives did not fully capture the target emotion only, and might have elicited also a mixture of emotions. Yet these may i) arise in different time points during the narratives and ii) be not as strong as the main target emotions (see Figure 1), thus average trial-wise activations most likely pertains to the target emotion. Despite this, the observed MVPA pattern may reflect whether each narrative is dominated by one emotion, or at least show the weighted influence of each emotion on a voxel activity. Therefore, the successful classification *per se* shows that at least the target emotions were successfully elicited, yet, better classification accuracies could potentially be reached if stimuli could be targeted more carefully to one category at the time.

## Conclusions

Our results characterize the discrete and distributed neural signatures of multiple emotional states. Different emotions result from differential activation patterns within a shared neural circuitry, mostly consisting of midline regions, motor areas, and subcortical regions. The more similar the neural underpinnings of these emotions, the more similarly they also are experienced. We suggest that the relative engagement of different parts of this system defines the current emotional state.

## Acknowledgements

We thank Marita Kattelus for her help in fMRI data collection and Matthew Hudson for a language check of the manuscript. Funding: This work was supported by the aivoAALTO project of the Aalto University, Academy of Finland (#265917 to L.N., and #138145 to I.P.J.), ERC Starting Grant (#313000 to L.N.); Finnish Cultural Foundation (#00140220 to H.S.); and the Swiss National Science Foundation National Center of Competence in Research for Affective Sciences (#51NF40-104897 to P.V.).

## Footnotes

1 For the sake of clarity, we use the term ‘non-basic emotions’ simply for emotions not belonging to the Ekman’s canonical list of basic emotions (anger, fear, disgust, happiness, sadness, surprise). In different emotion taxonomies, they could be called e.g. secondary or social.

